# Structure of a LRRC8 chimera with physiologically relevant properties reveals heptameric assembly and pore-blocking lipids

**DOI:** 10.1101/2022.07.28.501913

**Authors:** Hirohide Takahashi, Toshiki Yamada, Jerod S. Denton, Kevin Strange, Erkan Karakas

**Affiliations:** Department of Molecular Physiology and Biophysics, Vanderbilt University, School of Medicine, Nashville, TN, 37232, USA; Center for Structural Biology, Vanderbilt University; Nashville, TN, 37232, USA; Department of Anesthesiology, Vanderbilt University Medical Center, Nashville, TN, 37232, USA; Department of Pharmacology, Vanderbilt University, School of Medicine, Nashville, TN, 37232, USA

## Abstract

Volume-regulated anion channels (VRACs) mediate Cl^-^ and organic solute efflux from vertebrate cells and are essential for cell volume homeostasis. VRACs are heteromeric assemblies of LRRC8A-E proteins with unknown stoichiometries. Homomeric LRRC8A and LRRC8D channels have a hexameric structure. However, these channels are either non-functional or exhibit abnormal functional properties limiting their utility for structure-function analyses. We circumvented these limitations by developing novel homomeric LRRC8 chimeric channels with physiologically relevant functional properties. We demonstrate here that the LRRC8C-LRRC8A(IL1^25^) chimera comprising LRRC8C and 25 amino acids unique to the first intracellular loop (IL1) of LRRC8A has a heptameric structure like that of homologous pannexin channels. Membrane lipids are a key structural element of the channel and are located between subunits and occluding the channel pore. Our results suggest that native VRAC/LRRC8 channels are heptamers and that associated lipids are likely essential for normal channel gating and regulation.

## INTRODUCTION

<u>V</u>olume <u>R</u>egulated <u>A</u>nion <u>C</u>hannels, VRACs, are expressed widely in vertebrate cell types where they mediate the efflux of Cl^-^ and organic solutes required for cell volume regulation^1,2^. VRACs are activated by increases in cell volume and by large reductions in intracellular ionic strength^1^.

VRACs are encoded by the *Lrrc8* gene family^3,4^, which comprises five paralogs termed *Lrrc8a-e*^3,5^. Native VRAC/LRRC8 channels are heteromers with unknown stoichiometry. LRRC8A is an essential VRAC/LRRC8 subunit and must be co-expressed with at least one other paralog to reconstitute volume-regulated channel activity^3,6^. High resolution cryo-EM structures have been solved for homomeric LRRC8A^7–11^ and LRRC8D^12^ channels demonstrating that they have a hexameric configuration.

Defining the molecular basis by which VRAC and other volume-sensitive channels detect cell volume changes is a fundamental and longstanding physiological problem. Detailed molecular understanding of this important problem requires accurate channel structural information. Homomeric LRRC8A and LRRC8D channels have abnormal functional properties or are not expressed in the plasma cell membrane^3,13–15^. Structure-function studies of VRAC/LRRC8 heteromeric channels are complicated by their undefined, likely variable and experimentally uncontrollable stoichiometry. To circumvent these problems, we developed a series of novel homomeric LRRC8 chimeras that exhibit functional properties similar to those of native heteromeric VRAC/LRRC8 channels^14^.

We describe here the cryo-electron microscopy structure of the LRRC8C-LRRC8A(IL1^25^) chimera, hereafter termed 8C-8A(IL1^25^). 8C-8A(IL1^25^) consists of a 25-amino acid sequence unique to the first intracellular loop, IL1, of LRRC8A inserted into the corresponding region of LRRC8C (Supplementary Fig. 1a-c). Like native VRAC/LRRC8 channels, 8C-8A(IL1^25^) chimeras are activated strongly by cell swelling and low intracellular ionic strength, and the channels have normal pharmacological properties^13,14^. We demonstrate that the 8C-8A(IL1^25^) chimeric channel is a large-pore 7-subunit heptamer similar to homologous pannexin channels^16–23^. Membrane lipids are a key structural feature of 8C-8A(IL1^25^) and are located between channel subunits and occluding the channel pore, as has been shown recently for human pannexin 1^23^. Our results suggest that native VRAC/LRRC8 channels are heptamers and that associated membrane lipids likely play important roles in channel gating and regulation.

## RESULTS

### Structure determination

To facilitate structure-function understanding VRAC/LRRC8 channel regulation and pore properties, we expressed the 8C-8A(IL1^25^) chimera in Sf9 insect cells and purified the detergent-solubilized complexes by affinity and size exclusion chromatography (Supplementary Fig. 1). The purified 8C-8A(IL1^25^) complex appeared larger than the LRRC8A hexamers in both native-PAGE and size exclusion chromatography analysis (Supplementary Fig. 1e-f). We performed single-particle cryo-EM analysis to determine the structure of the complex (Supplementary Fig. 2-3). Following 2D classification and ab initio 3D reconstruction, we performed 3D classification and obtained five distinct 3D classes for 8C-8A(IL1^25^) (Supplementary Fig. 2). The 8C-8A(IL1^25^) maps showed no apparent symmetric arrangement. Therefore, the final reconstructions were done without enforcing any symmetry and resulted in five high-resolution (3.3-4.0 Å) structures (Figs. 1a,b; Supplementary Fig. 2-3).

**Figure 1:**
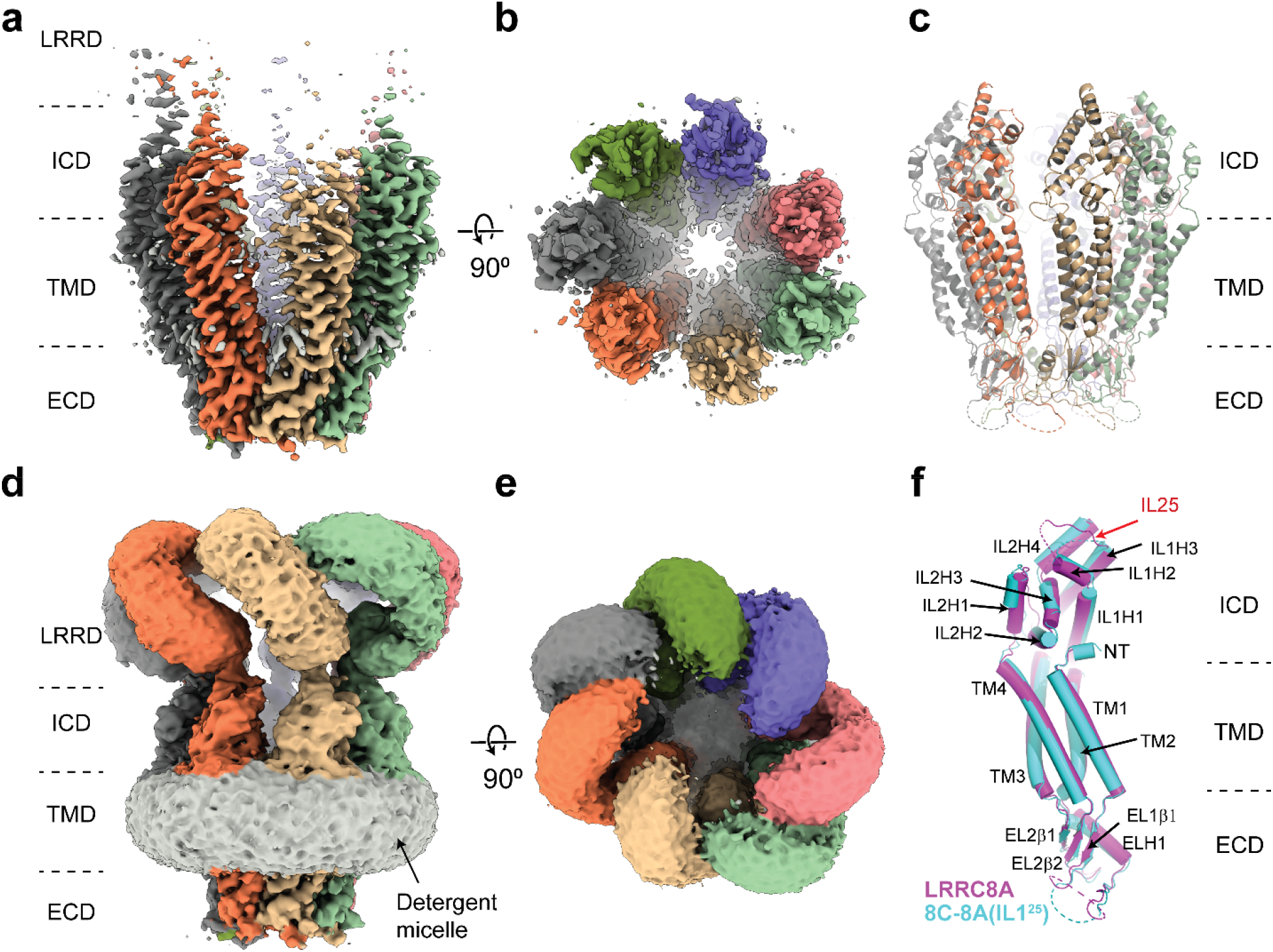
Cryo-EM structure of 8C-8A(IL1^25^) **a-b)** Cryo-EM maps of 8C-8A(IL1^25^) viewed through the membrane plane (**a**) and from the cytoplasm (**b**). **c**) Ribbon representation of the 8C-8A(IL1^25^) structure viewed through the membrane plane. **d-e)** Unsharpened Cryo-EM maps of 8C-8A(IL1^25^) viewed through the membrane plane (**d**) and from the cytoplasm (**e**), emphasizing low-resolution features. **f)** Structural comparison of the 8C-8A(IL1^25^) (cyan) and LRRC8A (magenta, PDB ID: 5ZSU) subunits.

Similar to LRRC8A and LRRC8D structures, the 8C-8A(IL1^25^) structure comprises four domains, the extracellular domain (ECD), transmembrane domain (TMD), intracellular domain (ICD), and leucine-rich repeat (LRR) motif-containing domain (LRRD) (Fig. 1). The cryo-EM maps for the ECD and TMD revealed high resolution features allowing us to build an atomic model comprising most of the protein except residues 60-94 in the ECD and the first 15 residues of the N-terminus. The quality of the cryo-EM maps for the ICDs was improved by performing local refinements, and the resulting maps were used to build a model (Supplementary Fig. 3). However, the first intracellular loop containing the swapped IL1^25^ region remained unresolved. Although the unsharpened maps revealed clear features for the entire protein, the local resolution for the LRRD was insufficient to build an atomic model, and the local refinement strategy we applied did not provide any meaningful improvement in LRRD resolution (Fig 1a-e; Supplementary Fig. 2-3). Therefore, we did not build an atomic model for the LRRD and used the maps without B-factor sharpening to assess their structure and overall arrangement relative to the rest of the protein complex (Fig. 1d-e).

### 8C-8A(IL1^25^) chimeras form heptameric channels

The overall structure of an individual 8C-8A(IL1^25^) protomer is similar to that of LRRC8A (Fig. 1f) and LRRC8D. However, unlike homomeric LRRC8A and LRRC8D channels, which are hexamers^7–12^, the 8C-8A(IL1^25^) chimeric channel is a 7 subunit heptamer, similar to the homologous pannexin channels (Figs. 1b and 1e; Supplementary Fig. 4) ^16–23^.

The subunit arrangement of the 8C-8A(IL1^25^) heptameric channel is asymmetric as opposed to LRRC8A and LRRC8D hexamers, which have 2-, 3-, or 6-fold symmetric arrangements^7–12^. When the ICD and TMD are viewed from the cytoplasm, 8C-8A(IL1^25^) protomers are organized into two distinct groups, one with four protomers and the other with three protomers (Fig. 2a-d). These two groups of protomers associate via a wide interface, while the protomers within each group associate via a narrow interface (Fig. 2a-d).

**Figure 2:**
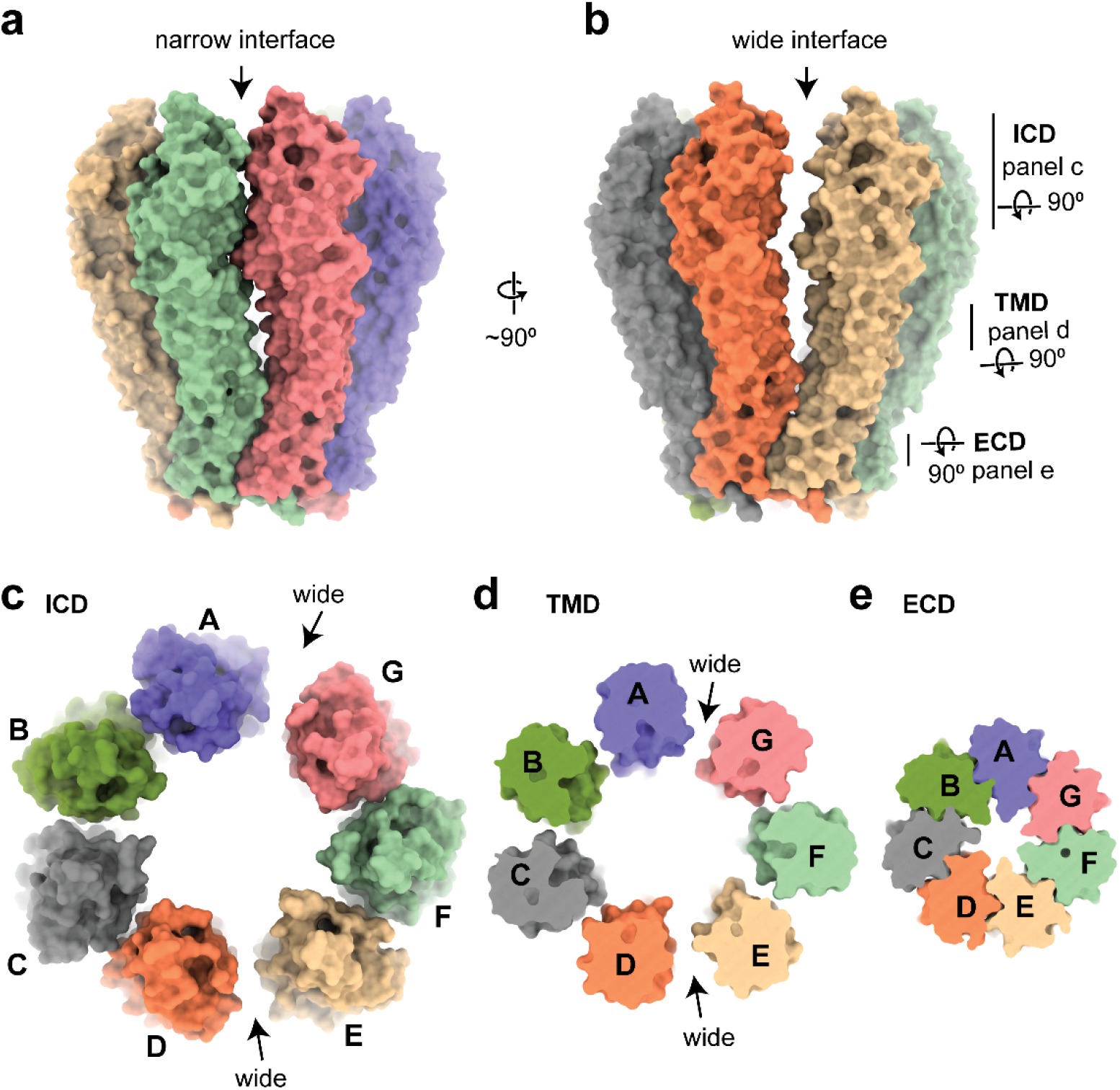
Subunit arrangement of the 8C-8A(IL1^25^) chimera. **a-b)** Surface representation of the 8C-8A(IL1^25^) chimera viewed from two sides, highlighting the “narrow” (**a**) and “wide” (**b**) interfaces. **c-e)** ICD, TMD, and ECDs are viewed from the cytoplasm with a depth of view as shown in panel **b**.

In contrast to the ICDs and TMDs, the ECDs are arranged symmetrically and form extensive contacts between the protomers (Fig. 2e). The asymmetric arrangement observed for the TMDs and ICDs is likely due to their divergent orientation relative to the ECDs. Figure 3a shows the alignment of the seven protomers on their ECDs. TMDs do not align and exhibit up to an 8° difference in their orientation relative to the ECD, while the difference is larger for the ICDs (Fig. 3a). The arrangement of the TMD and ICD relative to the ECD is also variable among different classes (Supplementary Fig. 5). Another notable difference between the protomers is in the linker that connects β-strand β1 of the ECD to the transmembrane helix 1 (TM1) of the TMD (Fig. 3a-b). In two of the protomers in the class 1 structure, K51 points toward the pore while D50 points in the opposite direction. In the five other protomers, D50 points toward the pore, and K51 is oriented toward the subunit interface (Fig. 3c). The distinct arrangement is also observed in other classes. However, the number of protomers with K51 pointing toward the pores differs. For example, the class 2 structure has only one protomer with K51 pointing toward the pore (Supplementary Fig. 6). It is plausible that this loop is flexible and adopts distinct conformations, possibly depending on the orientation of the TMD relative to the ECD.

**Figure 3:**
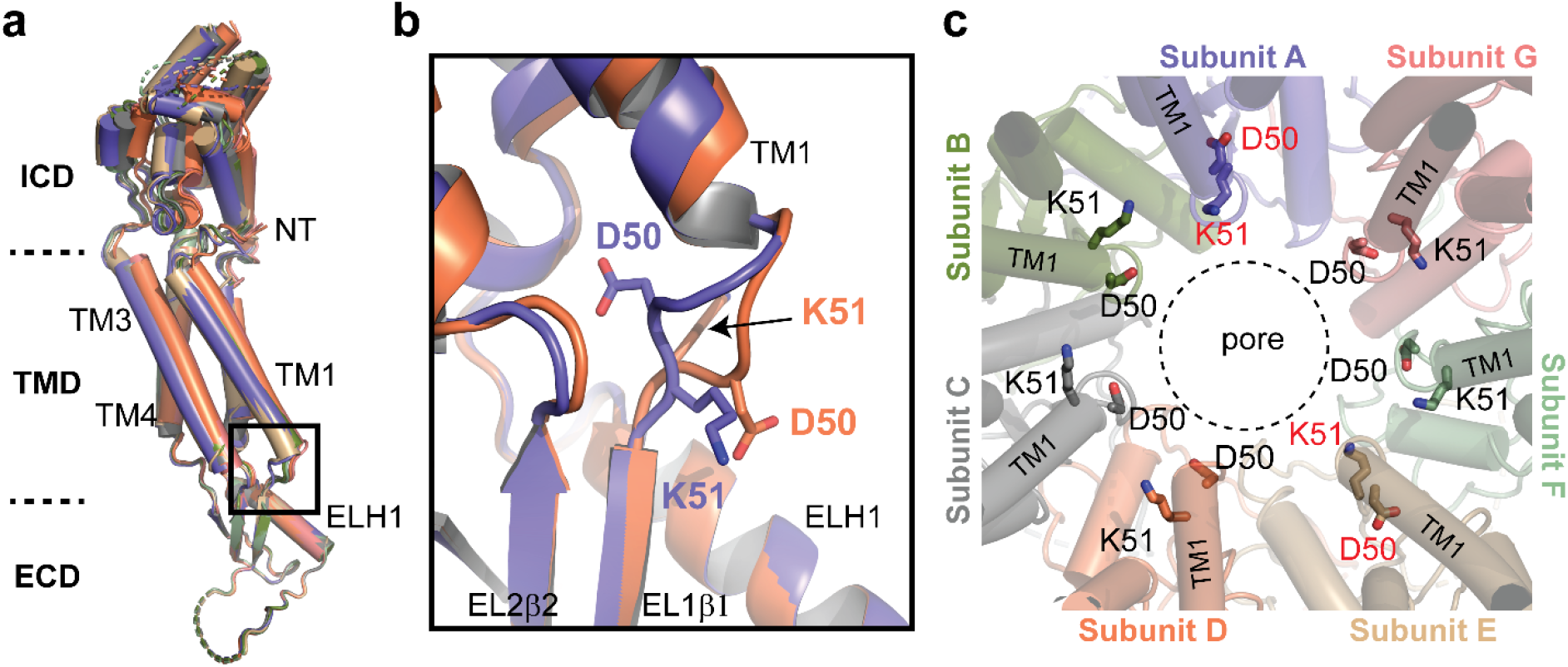
Structural heterogeneity of 8C-8A(IL1^25^) protomers. **a)** Structural comparison of the 8C-8A(IL1^25^) protomers (class 1). The structures are aligned based on their ECDs. **b**) Close-up view of the box region in panel **a**, highlighting the structural differences in the loop that connects TM1 to EL1β1. Only two protomers are shown. **c)** Close-up view of the pore around the residues D50 and K51, which are shown as sticks. The residues that adopt different conformation compared to others are labeled in red. The dashed circle indicates the pore-lining surface.

### Structural heterogeneity of 8C-8A(IL1^25^) chimeras

We obtained five distinct 3D classes for the 8C-8A(IL1^25^) chimera. The most apparent difference between the structures is in the arrangement of the LRRDs (Fig. 4a-b). In class 1, all seven LRRDs are arranged circularly with roughly 7-fold symmetry. For class 5, one of the LRRDs is positioned outside of the quaternary assembly formed by six LRRDs arranged with pseudo-2-fold symmetry. The density for the LRRD located outside the assembly is poorly visible, indicating high flexibility relative to the rest of the complex (Fig. 4a-b). The arrangements of the LRRDs in the other three 3D classes exhibit diverse arrangements and subunit interactions. As a result, several different subunit interfaces exist between the neighboring LRRDs. However, a detailed analysis of these interfaces is not possible due to the limited resolution of the cryo-EM maps for these regions, prohibiting the building of the models with amino acid assignments. Consistent with the conformational differences in the arrangements of the LRRDs, the ICDs also exhibit conformational heterogeneity, albeit less pronounced, among the five 3D classes we observed (Fig. 4c; Supplementary Fig. 5a-b).

**Figure 4:**
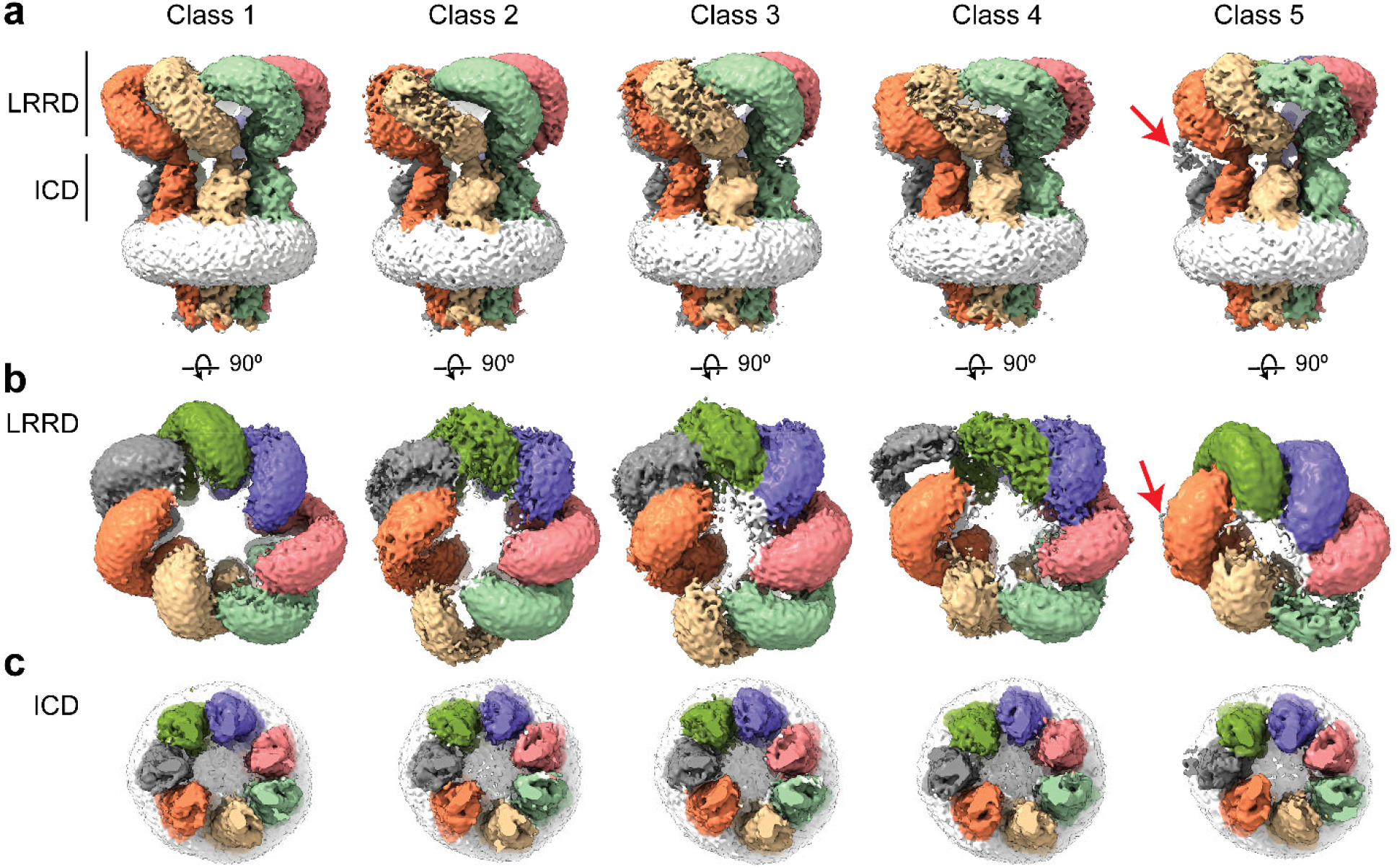
Conformational heterogeneity of 8C-8A(IL1^25^) chimeras. Cryo-EM maps (before sharpening) of 8C-8A(IL1^25^) 3D classes. Individual subunits are colored as in Fig 1. Detergent micelle is shown in white. Red arrows point to the weak density of the LRRD, which is not located within the LRRD quaternary assembly. **a**) Cryo-EM maps viewed through the membrane plane. Lines represent the depth of the view for LRRD and ICD. **b-c**). Cryo-EM maps viewed from cytoplasm for the LRRD (**b**) and ICD (**c**) with a depth as indicated in panel **a**.

We also observed structural differences in the TMDs among the different classes, but the overall clustered arrangement is preserved (Supplementary Fig. 5). Within the TMD, the positions of the protomers relative to the rest of the protein differ in different classes. This also affects the separation of the protomers from each other, especially at the wide interfaces. However, there are no apparent structural differences at the ECD of the structures (Supplementary Fig. 5c).

### Pore structure

Figure 5 shows the pore domain of the 8C-8A(IL1^25^) heptameric chimera compared to the pore domain of LRRC8A hexameric channels. The narrowest region of the heptameric 8C-8A(IL1^25^) channel pore has an average solvent-accessible radius of 4.7 Å and is formed by L105 located on the extracellular side of the protein (Fig. 5a). This is considerably larger than the pore radius of 2.0 Å for hexameric LRRC8A channels (Fig. 5b-c)^7–10^.

**Figure 5:**
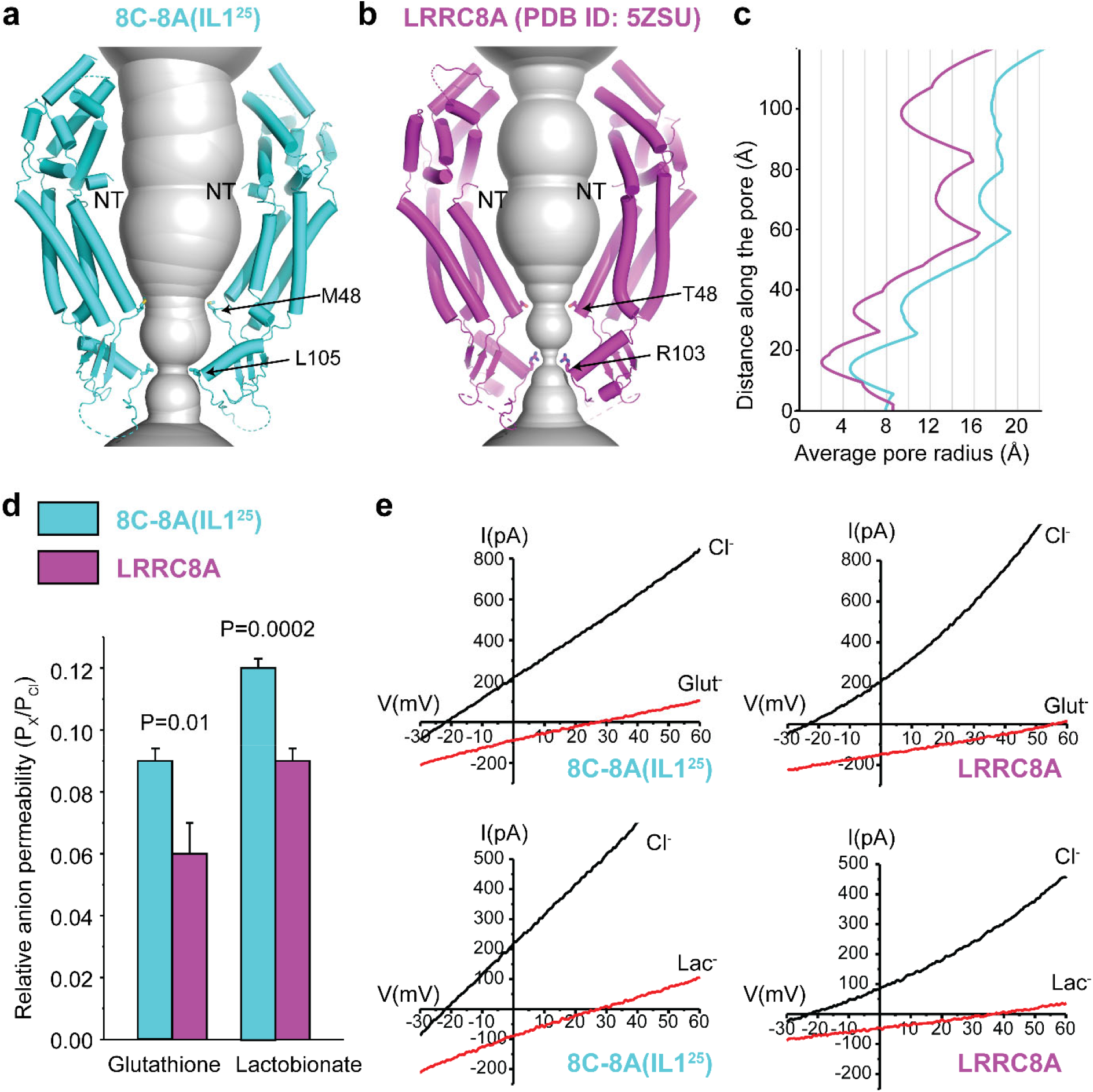
Comparison of channel pores. Pore openings of the 8C-8A(IL1^25^) heptameric channel (**a**) and 8A (PDB ID: 5ZSU) homohexameric channel (**b**) calculated using the software program HOLE^61^. Only two opposing subunits are shown. Residues forming the constriction sites are shown as the sticks. The first modeled residues at the N-termini are labeled as NT. **c**) 1D graph of the average radius along the length of the 8C-8A(IL1^25^) (cyan) and 8A (purple) channel pores. **d**) Relative (P_x_/P_Cl_) glutathione and lactobionate permeabilities calculated from reversal potential changes induced by replacing bath Cl^-^ with the test anion. Values are means ± SEM (N=4-7). **e**) Representative LRRC8A and 8C-8A(IL1^25^) current traces in the presence of bath Cl^-^ or after substitution with glutathione (Glut^-^) or lactobionate (Lac^-^). Currents were elicited by ramping membrane voltage from −100 mV to +100 mV.

To further assess the pore diameters of the two channels, we quantified the relative permeabilities (i.e., P_x_/P_Cl_) of LRRC8A and 8C-8A(IL1^25^) to the large organic anions glutathione and lactobionate. Consistent with its cryo-EM structure, P_glutathione_/P_Cl_ and P_lactobionate_/P_Cl_ were both significantly (P<0.01) higher for the 8C-8A(IL1^25^) chimeric channel (Fig. 5d-e).

### Interaction of lipids with the 8C-8A(IL1^25^)

As shown in Fig. 6a, we observe non-protein densities penetrating through the openings between the protomers in the TMD. The shape and length of these densities are consistent with lipid molecules. Similar densities are observed in LRRC8A hexameric channels reconstituted in lipid nanodiscs^7^. However, unlike the LRRC8A hexameric channel structures^7^, we also observed two layers of thick densities that run parallel to the membrane plane within the pore (Fig. 6b). These densities are aligned with the boundaries of the detergent micelle and plausibly represent the lipid bilayer obstructing the pore as suggested for other large-pore channels, including pannexins and innexins^23,24^. The lipid bilayer blocks the entire pore, making the channel impermeable to solutes. Hence, the structures presented here likely represent closed channel conformations.

**Figure 6.**
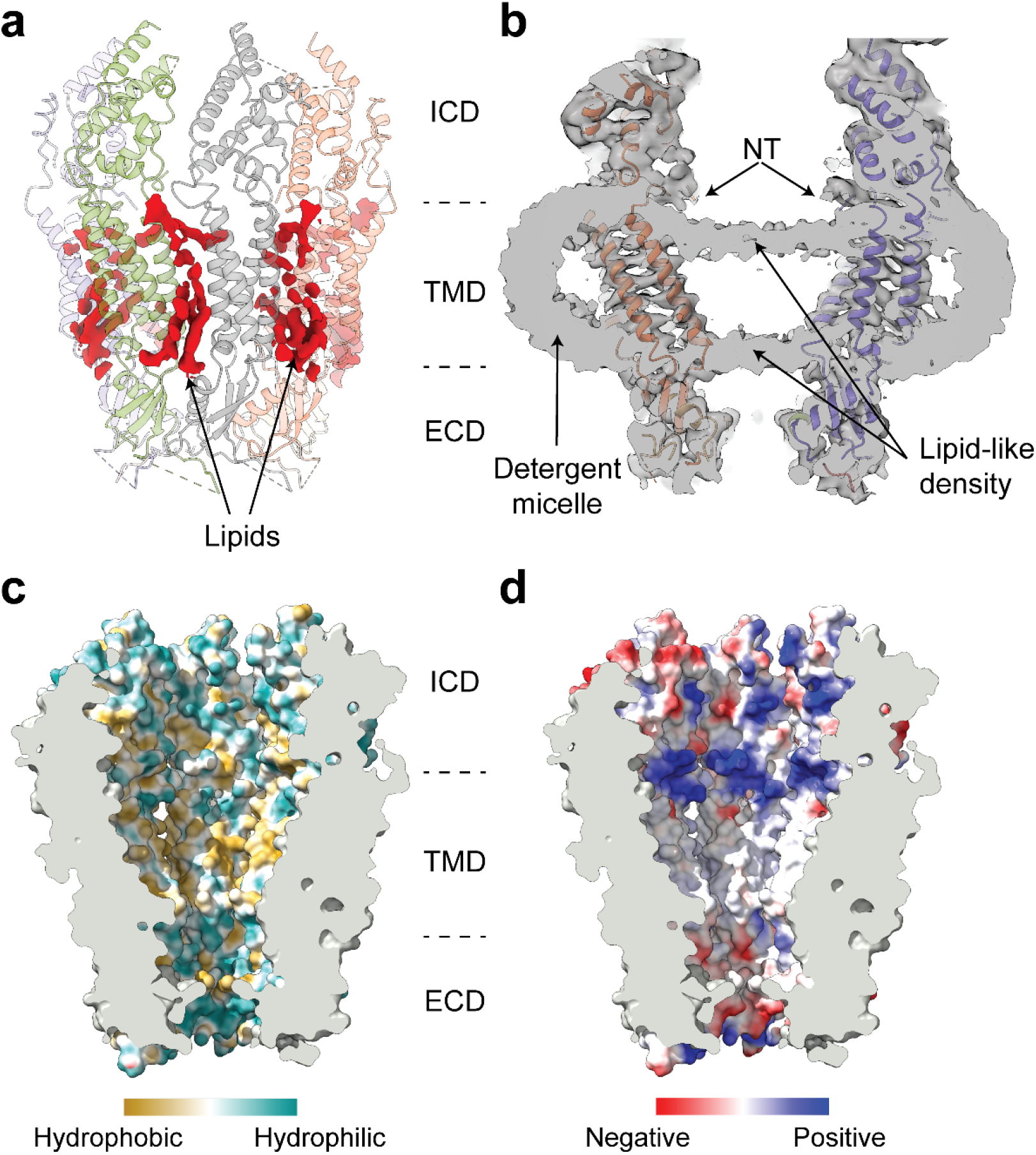
Interaction of lipids with the 8C-8A(IL1^25^) chimera. **a**) Ribbon representation of the 8C-8A(IL1^25^) channel (transparent) along with the cryo-EM densities (red) of the lipid molecules between the subunits viewed through the membrane plane. **b**) A sliced view of the unsharpened cryo-EM map (grey, transparent) with the ribbon representation of the 8C-8A(IL1^25^) structure. Arrows indicate densities corresponding to pore-blocking lipids and detergent micelles. **c-d)** Surface representation of the 8C-8A(IL1^25^) pore colored based on hydrophobicity (**c**) and electrostatic charge (**d**).

To confirm that these densities are not detergent-induced artifacts, we reconstituted the 8C-8A(IL1^25^) chimera in lipid nanodiscs and performed cryo-EM analysis (Supplementary. Fig 7). Although the resolution of the cryo-EM maps could not be improved beyond 7 Å, the same bilayer-like density within the channel pore was visible (Supplementary Fig. 7).

Between the two lipid layers, the inner surface of the pore is primarily hydrophobic, creating a favorable environment for the acyl chains of the phospholipids (Fig. 6c). The bilayer leaflet facing toward the intracellular space is surrounded by positively charged residues located underneath the N-terminal region of the protein (Fig. 6d). The bilayer leaflet facing towards the extracellular space is encircled by M48 located at the C-terminal end of TM1. Both leaflets are aligned with the lipids penetrating through the interfaces between the protomers, suggesting a plausible route for the lipids to move in and out of the pore (Fig. 6).

## DISCUSSION

LRRC8A and LRRC8D cryo-EM structures provided new insights into LRRC8 protein secondary and tertiary structure and channel quaternary conformational arrangements. However, utilizing protein structure to gain detailed molecular insights into protein function and regulation requires targeted mutagenesis and functional analysis guided by accurate structural details.

With the exception of LRRC8A, homomeric LRRC8 channels do not traffic to the plasma membrane and cannot be functionally characterized^3,14,15^. LRRC8A homomers form plasma membrane channels, but they exhibit abnormal functional properties. Most notably, LRRC8A channels are not activated by cell swelling under normal physiological conditions^13,14^ and are only weakly activated by extreme reductions in cytoplasmic ionic strength (Supplementary Figure 8)^13^. LRRC8A channels also exhibit abnormal pharmacology^13^. The non-native functional properties of LRRC8A channels indicate that they have a non-native structure, which presents significant limitations for understanding how VRAC/LRRC8 channels are regulated and how pore structure determines solute permeability characteristics.

Homomeric LRRC8 channels with physiologically relevant functional properties are formed by chimeric proteins containing parts of the essential subunit LRRC8A and another LRRC8 paralog^14^. The 8C-8A(IL1^25^) chimeric channel exhibits normal sensitivity to cell volume changes and intracellular ionic strength and normal pharmacology (Supplementary Figure 8)^13,14^.

Unlike homomeric LRRC8A and LRRC8D channels, which are hexamers, our studies demonstrate that 8C-8A(IL1^25^) chimeric channels have a heptameric structure (Figs. 1–2). The number of subunits required to form native VRAC/LRRC8 channels is unknown. However, native gel, crosslinking and mass spectrometry studies^6^ and photobleaching experiments^25^ suggest that VRAC/LRRC8 channels are formed by >6 LRRC8 subunits.

The 8C-8A(IL1^25^) has a limiting pore radius of 4.7 Å (Figs. 4a and 4c; Supplementary Fig. 9a), which unlike the much narrower LRRC8A pore (Fig. 4b and 4c; Supplementary Fig. 9b), is very consistent with the 6-7 Å pore radius estimated for native VRAC/LRRC8 channels^26,27^ and the role of these channels in the transport of large anionic and uncharged organic solutes^1,28^. A leucine residue located at position 105 forms the 8C-8A(IL1^25^) pore constriction (Fig. 4b). In LRRC8A and LRRC8B, L105 is replaced by arginine. LRRC8D and LRRC8E paralogs have either phenylalanine or leucine at the homologous position. Hexameric LRRC8A and LRRC8D channels have limiting pore radii of 2.0 Å and 3.5 Å, respectively (Fig. 4b; Supplementary Fig. 9). When we create hypothetical heptamers by aligning LRRC8A and LRRC8D subunits onto individual 8C-8A(IL1^25^) protomers, the pore radii increase to 4.7 Å and 7.1 Å, respectively, indicating that the oligomeric state has a direct impact in pore size and likely in channel transport properties (Supplementary Fig. 9d-f).

The abnormally narrow pore radius of 2 Å of the LRRC8A hexameric channel (Fig. 4b) likely accounts for its abnormal DCPIB pharmacology. Native VRAC/LRRC8 channels are inhibited >90% by 10 μM DCPIB. Inhibition is voltage-insensitive and exhibits a Hill coefficient of 2.9^29^ suggesting that multiple DCPIB molecules are required to inhibit the channel. The DCPIB pharmacology of 8C-8A(IL1^25^) fully recapitulates that of native VRAC/LRRC8 channels^13^. In contrast, LRRC8A hexameric channels are weakly inhibited by 10 μM DCPIB, and inhibition is strongly voltage-dependent with a Hill coefficient close to 1^13^. The low Hill coefficient is consistent with a LRRC8A cryo-EM structure described by Kern et al.^7^ showing a single DCPIB molecule blocking the channel pore at is the narrowest constriction formed by R103 (Fig 5b).

LRRC8 proteins are most closely related to pannexin channel-forming proteins and appear to have diverged from this gene family at the origin of the chordates^5^. Low resolution structural studies led to the early incorrect conclusion that pannexins form hexameric channels^30–32^. However, several recent high resolution cryo-EM studies have demonstrated that pannexin channels have a 7-subunit, heptameric structure^16–23^.

Our findings taken together with biochemical and fluorescence photobleaching measurements^6,25^ and the well-defined heptameric structure of closely related pannexin channels^16–23^ suggest that native VRAC/LRRC8 channels are likely heptamers formed by LRRC8A and at least one other LRRC8 paralog. However, it is conceivable that native channels exist as both hexamers and heptamers and possibly higher order oligomers as has been reported for CALHM channels^33^. Defining the stoichiometry and subunit arrangement of native VRAC/LRRC8 channels remains an important challenge in the field.

Our studies revealed that lipids are key structural components of the 8C-8A(IL1^25^) chimeric channel. Kern et al. ^7^ identified lipids located between individual subunits in LRRC8A hexameric channels reconstituted in lipid nanodiscs. Intersubunit non-protein densities, which are most likely lipids, are also observed between 8C-8A(IL1^25^) subunits in detergent reconstituted channels (Fig. 6A). Importantly, we identified for the first time two layers of lipids blocking the pore of the 8C-8A(IL1^25^) heptameric channel (Fig. 6b; Supplementary Fig. 7). Intrapore lipids have recently been shown to be critical structural and regulatory elements of pannexins^23^ and closely related innexin^24^ CALHM^34,35^ channels, as well as bacterial mechanosensitive channels^36–38^.

Similar to what we observe with the 8C-8A(IL1^25^) heptameric channel, Kuzuya et al. ^23^ recently demonstrated the presence of two layers of lipids blocking the pore of the human pannexin-1 channel. Their studies suggest that movement of these lipids out of the pore through gaps between the subunits is required for channel opening and that lipid migration is regulated by an α-helix located on the short N-terminus of the pannexin-1 protein. Location of the N-terminal α-helices in the pore may block lipid entry and maintain the channel in an open state, while closure may be mediated by the interaction of the N-terminal α-helix with the cytoplasmic C-terminus allowing lipids to enter the pore.

It is noteworthy that the short N-terminus of LRRC8 proteins contains a coiled domain that was identified as an α-helix in LRRC8D^12^. The location of the N-terminus is poorly resolved in most cryo-EM structures, including that of the 8C-8A(IL1^25^) heptameric channel. However, in LRRC8D cryo-EM structures, the channel N-terminus is resolved and shows the helix protruding into the pore^12^. One human LRRC8A cryo-EM structure shows that the N-terminus is highly coordinated with the cytoplasmic C-terminus^9^. Furthermore, mutagenesis and electrophysiology studies suggest that the N-terminus plays a role in channel gating and in controlling pore properties^9,15^. Given the close evolutionary relationship between pannexin and LRRC8 proteins, and the striking structural similarities of 8C-8A(IL1^25^) and pannexin channels, it is reasonable to postulate that VRAC/LRRC8 channels are gated by the pore lipids we identified functioning together with the N-terminus.

Like VRAC/LRRC8 channels, mechanosensitive channels mediate the osmoregulatory efflux of solutes in bacteria following hypo-osmotic shock^39^. The small conductance bacterial mechanosensitive channel MscS is a homoheptamer activated by increases in membrane bilayer tension. Lipids are key structural elements of these channels and are located in the channel pore and between subunits^36–38^. Movement of lipids out of the pore gate the channel open^38^. Interestingly, MscS channels enter a non-conducting state that cannot be activated even with increased membrane tension. Zhang et al.^38^ proposed that loss of closely associated lipids gives rise to this desensitized, non-conducting channel state. By analogy, the poor sensitivity of LRRC8A homomeric channels to cell swelling and reduced intracellular ionic strength^13,14^ (Supplementary Figure 8) may be due to the absence of properly associated lipids.

Leucine-rich repeat motifs have been identified in over 14,000 proteins from viruses to eukaryotes and play essential roles in signal transduction and as sites for protein-protein interactions^40–42^. The LRR motif can also function as a mechanosensor^43,44^. VRAC/LRRC8 channels most likely arose by fusion of the transmembrane domain encoding portion of a pannexin channel gene with the LRR domain portion of an unrelated gene type^5^. Conformational changes in the VRAC/LRRC8 channel LRR domain are correlated with changes in channel activity^11,45^ and multiple cryo-EM structures demonstrate that this region is conformationally flexible (Fig. 3) ^7–12^. These data all point to the high likelihood that the VRAC/LRRC8 channel LRR domain is a critical element of the channel’s cell volume sensing apparatus.

LRRC8 chimeras with normal cell volume and ionic strength sensitivity must contain all or part of the LRRC8A IL1^14^ indicating that this protein region is also critical for channel regulation. The 25 amino acid region of the LRRC8A IL1 inserted into the 8C-8A(IL1^25^) chimera is predicted to be intrinsically disordered^14^. Consistent with this prediction, this region was not visible in our cryo-EM structures. It is conceivable that the LRRC8A IL1 is required for correct channel assembly and conformation, that it plays a direct role in cell volume sensing and/or that it functions to transduce cell volume-induced conformational changes into changes in the gating.

The mechanisms by which eukaryotic cells sense cell volume changes and transduce those changes into regulatory responses remain mysterious. Understanding how VRAC/LRRC8 channels detect cell volume is greatly constrained by the abnormal functional properties of LRRC8A homomers and by the undefined heteromeric nature of native channels. The high resolution cryo-EM structures of the 8C-8A(IL1^25^) homomeric chimera, which has native physiological properties, has revealed several novel structural elements of VRAC/LRRC8 channels. These structures now provide the foundation for defining the molecular basis of channel cell volume sensing utilizing structure-guided mutagenesis combined with electrophysiological functional analysis of channels with defined stoichiometry and subunit arrangement.

## METHODS

### Constructs

Human LRRC8A and LRRC8C cDNAs cloned into pCMV6 were purchased from OriGene Technologies. The 8C-8A(IL1^25^) chimera cDNA construct was generated using the Phusion High-Fidelity PCR kit (New England BioLabs). All cDNAs were tagged on their carboxy terminus with Myc-DDK epitopes. For protein expression, the cDNAs encoding human LRRC8A and the 8C-8A(IL1^25^) chimera were sub-cloned with a C-terminal FLAG tag into pAceBac1 vectors and incorporated into baculovirus using the Multibac expression system^46^. All constructs were verified by DNA sequencing.

### Protein expression and purification

8C-8A(IL1^25^) chimera was expressed in *Sf9* cells (4×10^6^ cells/ml) at 27 °C for 48 hours. Cells were harvested by centrifugation (2,000*g*) and resuspended in a lysis buffer composed of 150mM NaCl and 50 mM Tris-HCl, pH 8.0, 1 mM Phenylmethylulfonyl fluoride (PMSF). After cell lysis using Avastin EmulsiFlex-C3, the cell lysate was clarified from large debris by centrifugation at 6,000*g* for 20 min. The cleared lysate was centrifuged at 185,000 x *g* (Type Ti45 rotor) for 1 h. Membrane pellets were resuspended and homogenized in ice-cold resuspension buffer (150 mM NaCl, 50 mM Tris-HCl, pH 8.0) and solubilized using 0.5% lauryl maltose neopentyl glycol (LMNG) at a membrane concentration of 100 mg/ml. The solubilized pellets were stirred gently for 4 h at 4 °C, and the insoluble material was separated by centrifugation at 185,000 x *g* (Type Ti45 rotor) for 40 min. The supernatant was then mixed with anti-FLAG affinity gel resin (Sigma) at 4 °C for 1 hour. After washing the resin with 10 column volume wash buffer composed of 150 mM NaCl, 50 mM Tris-HCl, pH 8.0, and 0.005% LMNG, the protein was eluted using the wash buffer supplemented with 100μg/ml FLAG peptide. Protein was further purified by size exclusion chromatography using a Superose 6 Increase column (10/300 GL, GE Healthcare) equilibrated with 150 mM NaCl, 50 mM Tris-HCl, pH 8.0, 0.005% LMN. The peak fraction corresponding to 8C-8A(IL1^25^) was concentrated to 3.0 mg/ml, centrifuged at 70,000 rpm using an S110-AT rotor (Thermo Scientific) for 10 min, and used immediately for cryo-EM imaging. Human LRRC8A with a C-terminal Flag tag was purified using the same protocol described above for 8C-8A(IL1^25^).

For nanodisc incorporation, 8C-8A(IL1^25^) chimera was purified as described above except n-Dodecyl-β-D-Maltopyranoside (DDM, 1% for solubilization of the membrane and 0.05% in the purification buffers) was used instead of LMNG. The membrane scaffold protein MSP1E3D1 was expressed using the p MSP1E3D1 plasmid, a gift from Stephen Sligar (Addgene plasmid # 20066; http://n2t.net/addgene:20066; RRID: Addgene_20066)^47^. MSP1E3D1 was expressed in *E. coli* BL21(DE3) cells and purified as described previously using Ni^2+^ affinity resin^48^. The histidine tag was cleaved off using TEV protease, and the protein was further purified by SEC using a HiLoad 16/600 Superdex 200 pg column equilibrated with 300 mM NaCl and 40 mM Tris-HCl, pH 8.0. Peak fractions containing MSP1E3D1 were collected and stored at −80°C for the nanodisc formation.

### MSP1E3D1 Nanodisc formation

The preparation of nanodiscs was performed by mixing 8C-8A(IL1^25^) purified in the presence of DDM with POPC lipids (Avanti Polar Lipids, Inc.) and MSP1E3D1 at a final molar ratio of 1:2.5:250 (8C-8A(IL1^25^):MSP1E3D1:POPC). The mixture was incubated with Biobeads SM2 (Bio-Rad) overnight. After removing the biobeads by centrifugation, the protein sample was concentrated and purified by SEC using a Superose 6 10/300 Increase column equilibrated with 150 mM NaCl and 50 mM Tris-HCl, pH 8.0. Peak fractions corresponding to 8C-8A(IL1^25^)- MSP1E3D1 nanodiscs were concentrated to 2.0 mg/ml and used for cryoEM grid preparation immediately.

### Cryo-EM sample preparation and data collection

Purified 8C-8A(IL1^25^) was applied to 300 mesh UltrAuFoil holey gold 1.2/1.3 grids (Quantifoil Microtools) that were glow discharged for 10 seconds at 25 mA. The grids were blotted for 4 seconds at force 12 using double-layer Whatman filter papers (1442-005, GE Healthcare) before plunging into liquid ethane using an FEI MarkIV Vitrobot at 8 °C and 100% humidity. Samples were imaged using a 300 kV FEI Krios G3i microscope equipped with a Gatan K3 direct electron camera. Movies containing 40 frames were collected in super-resolution mode at 81,000x magnification with a physical pixel size of 1.1 Å/pixel and defocus values at a range of −0.8 to −1.5 μm using the automated imaging software SerialEM^49^.

8C-8A(IL1^25^) nanodisc grids were prepared as described above. Samples were imaged using a 300 kV FEI Krios G4 microscope equipped with a Gatan K3 direct electron camera. Movies containing 50 frames were collected in super-resolution mode at 103,000x magnification with a physical pixel size of 0.818 Å/pixel and defocus values at a range of −0.8 to −2.2 μm using the automated imaging software EPU (ThermoFisher Scientific).

### Cryo-EM data processing

All image processing was performed using CryoSparc2^50^. Motion correction and CTF estimations were performed locally using Patch Motion Correction and Patch CTF Estimation procedures. Initial particle picking was performed by blob search. Particles were then binned 4x and extracted. After 2D classification, classes with clear structural features were selected and used for template-based particle picking. The new set of particles was binned four times and extracted. After 2D classification, a cleaned particle set was used for the ab initio 3D reconstruction. The resulting map was used as a model for 3D classification. After a series of 2D and 3D classification runs, particles were reextracted with a box size of 360×360 pixels. These particles were separated into six classes using 3D classification. Five out of six classes revealed interpretable density maps, and these particles were processed further using nonuniform refinement to obtain the final cryo-EM maps for each class.

To improve the quality of the cryo-EM maps around the ICDs, we performed local refinements using masks that cover the ICD of each protomer (Supplementary Fig. 3). We observed a noticeable improvement in class 1, 2, and 3 structures and used these maps for model building. We applied the same strategy for the LRRDs. However, the quality of the maps did not improve to a level allowing us reliable model building.

Data processing for 8C-8A(IL1^25^) nanodiscs was performed as described above, with the following exceptions. Motion correction was performed using MotionCor2^51^ in Relion 3.0^52^. The final set of particles was extracted with a box size of 480×480 pixels, and they were classified into three 3D classes.

### Model building

Models were built using Coot^53^. We initially placed the human LRRC8C model from the AlphaFold protein structure prediction database^54,55^ in the class 1 density. We manually fitted the individual residues into the density while removing the parts that do not have corresponding interpretable densities. Once we built a complete protomer, we copied the model into the other six protomers and manually fit individual residues into the density. The model was refined using Phenix real-space refinement^56^. We performed iterative build-refine cycles till a satisfactory model was obtained. The resulting model was fitted into the class 2 and 3 maps and fitted into the density using the same build-refine iterations as described above. Because of their limited resolution, most parts of class 5 and 6 structures were modeled without their side chains (i.e., as alanines) while maintaining their correct labeling for the amino acid type. The LRRDs were not modeled in all five structures. Validations of the structural models were performed using MolProbity^57^ implemented in Phenix^56^.

Some of the data processing and refinement software was supported by SBGrid^58^.

### Patch clamp electrophysiology

LRRC8A and 8C-8A(IL1^25^) cDNA constructs were expressed in *Lrrc8*^-/-^ human colon cancer HCT116 cells in which the five *Lrrc8* genes were disrupted by genome editing. *Lrrc8*^-/-^ cells were a generous gift from T. Jentsch. Cells were transfected using Turbofectin 8.0 (OriGene Technologies) with 0.125 μg GFP cDNA and 0.25 μg of 8A and 8C-8A(IL1^25^). All experimental protocols were performed on at least two independently transfected groups of cells.

Transfected cells were identified by GFP fluorescence and patch clamped in the whole cell mode at room temperature using patch electrodes pulled from 1.5 mm outer-diameter silanized borosilicate microhematocrit tubes. Recordings were not performed on cells where access resistance was >2-fold that of the pipette resistance.

The composition of the control bath and pipette solutions used in these studies is shown in Table 1. Intracellular ionic strength was reduced by the removal of CsCl or Cesium methanesulfonate from the patch pipette solution and replacement with sucrose to maintain solution osmolality.

**Table 1.** Composition of patch pipette and solutions.

|  | **Patch pipette solutions** |  | **Bath solutions** |  |
| --- | --- | --- | --- | --- |
|  | **Control** | **Control** | **Control** | **Hypotonic** |
| CsCl | 126 mM | 26 mM | 75 mM | 75 mM |
| Cs methanesulfonate |  | 100 mM |  |  |
| MgSO<sub>4</sub> | 2 mM | 2 mM | 5 mM | 5 mM |
| Ca-gluconate<sub>2</sub> |  |  | 1 mM | 1 mM |
| ATP-Na<sub>2</sub> | 2 mM | 2 mM |  |  |
| GTP-Na<sub>2</sub> | 0.5 mM | 0.5 mM |  |  |
| Glutamine |  |  | 2 mM | 2 mM |
| EGTA | 1 mM | 1 mM |  |  |
| HEPES | 20 mM | 20 mM | 12 mM | 12 mM |
| Tris |  |  | 8 mM | 8 mM |
| CsOH | 12 mM | 12 mM |  |  |
| HCl |  |  | 2 mM | 2 mM |
| Glucose |  |  | 5 mM | 5 mM |
| Sucrose | 16 mM | 16 | 115 mM | 70 mM |
| pH | 7.2 | 7.2 | 7.4 | 7.4 |
| Osmolality | 275 mOsm | 275 mOsm | 300 mOsm | 250 mOsm |
| Ionic strength | 0.162 M | 0.162 M |  |  |

The pH of patch pipette and bath solutions was adjusted with CsOH and HCl, respectively.

Whole-cell currents were measured with an Axopatch 200A (Axon Instruments) patchclamp amplifier. Electrical connections to the patch-clamp amplifier were made using Ag/AgCl wires and 3M KCl/agar bridges. Series resistance was compensated by >85% to minimize voltage errors. Data acquisition and analysis were performed using pClamp 10 software (Axon Instruments).

Changes in current amplitude were quantified using a voltage-ramping protocol. Membrane voltage was held at −30mV throughout all experiments. Ramps were initiated by stepping membrane voltage to −100mV and then ramping membrane voltage over 1 s to +100mV. This was followed by a step back to −30mV for 4 s before the ramp was repeated.

Relative anion permeability (P_x_/P_Cl_) was measured from Cl^-^ substitute induced changes in reversal potential using a modified Goldman–Hodgkin–Katz equation^3^. LRRC8A and 8C-8A(IL1^25^) currents were activated with a low ionic strength, 0.062 M, patch pipette solution and by cell swelling induced by reducing bath osmolality to 250 mOsm. After stable current activation was achieved, changes in reversal potential were induced by replacing bath CsCl with either cesium glutathione or cesium lactobionate. Reversal potentials were corrected for anion-induced changes in liquid junction potentials.

## Figure preparation

Figures were prepared using Chimera^59^, ChimeraX^60^, and The PyMOL Molecular Graphics System (Version 2.0, Schrödinger, LLC). Calculation of the pore radii was performed using the software HOLE^61^.

## Acknowledgments

We thank Dr. Kunpeng Lee at Case Western Reserve University and Melissa Chambers and Scott Collier at the Cryo-EM facility at Vanderbilt University for cryo-EM data collection. This work was conducted in part using the CPU and GPU resources of the Advanced Computing Center for Research and Education (ACCRE) at Vanderbilt University. We used the DORS storage system supported by the National Institute of Health (NIH) (S10 RR031634 to Jarrod Smith). This work was supported by the National Institute of Diabetes, Digestive, and Kidney Diseases Grant R01 DK51610 to JSD.

## Author contributions

Conceptualization- JSD, EK, KS; methodology- HT, TY; investigation- HT, TY; formal analysis- JSD, EK, KS, HT, TY; writing- original draft- EK, KS; writing- review and editing- JSD, EK, KS, HT, TY; supervision- EK, JSD, KS; project administration- EK, JSD, KS; funding acquisition- EK, JSD, KS.

