## Supplementary Material for "Structure of a LRRC8 chimera with physiologically relevant properties reveals heptameric assembly and pore-blocking lipids"

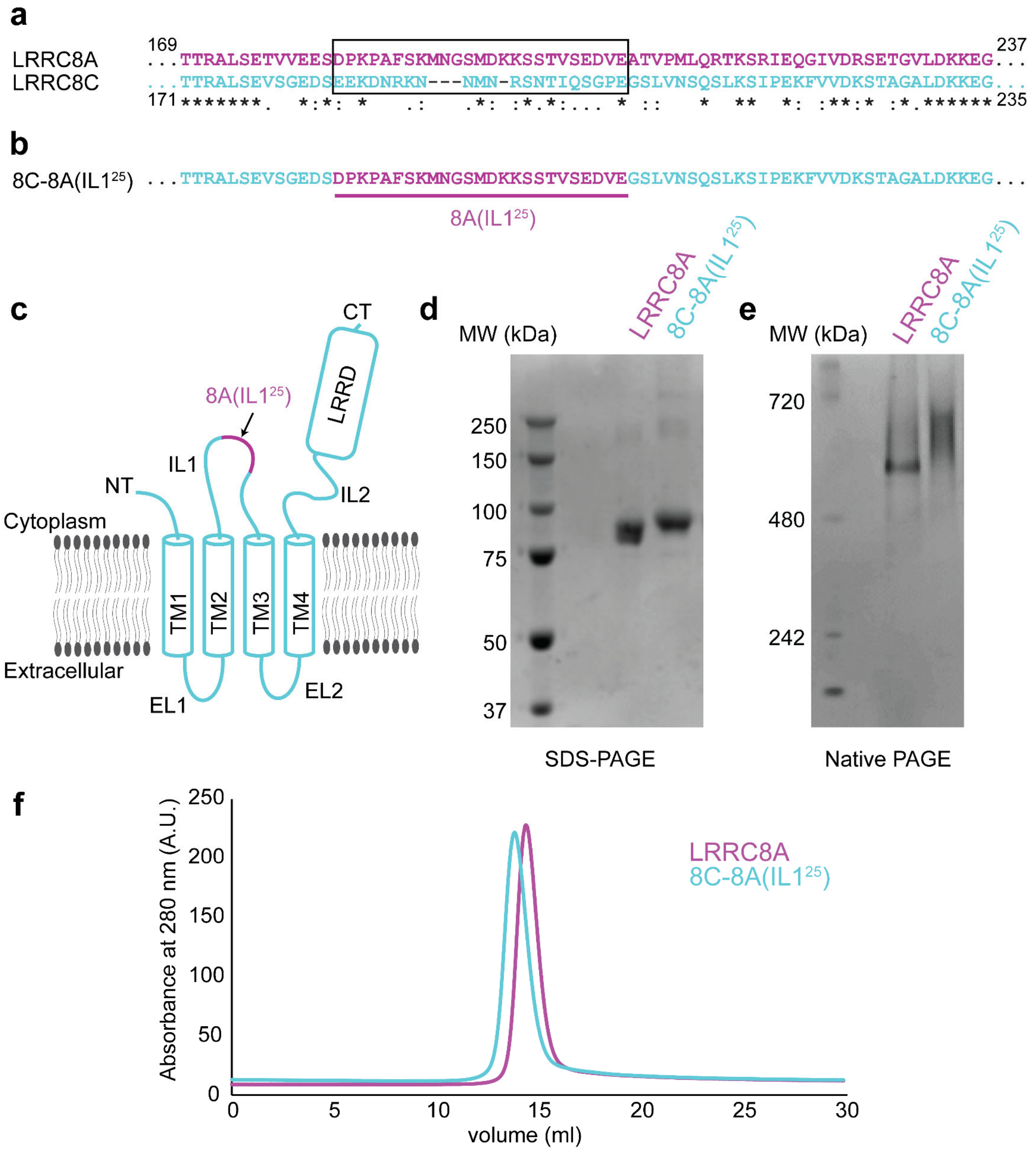


**Supplementary Fig. 1: Construct design and purification of 8C-8A(IL1$^{25}$). a)** Sequence alignment of LRRC8A and LRRC8C around the swapped IL1$^{25}$ region. The region that is swapped in the 8C-8A(IL1$^{25}$) construct is shown inside a black box. **b)** The amino acid sequence of the 8C-8A(IL1$^{25}$) construct around the swapped IL1$^{25}$ region. **c)** Schematic diagram of the 8C-8A(IL1$^{25}$) protein highlighting the relative position of the swapped IL1$^{25}$ region. **d-f)** SDS-PAGE (**d**) and native PAGE (**e**) and size exclusion chromatography (**f**) analysis of the purified LRRC8A and 8C-8A(IL1$^{25}$) proteins.

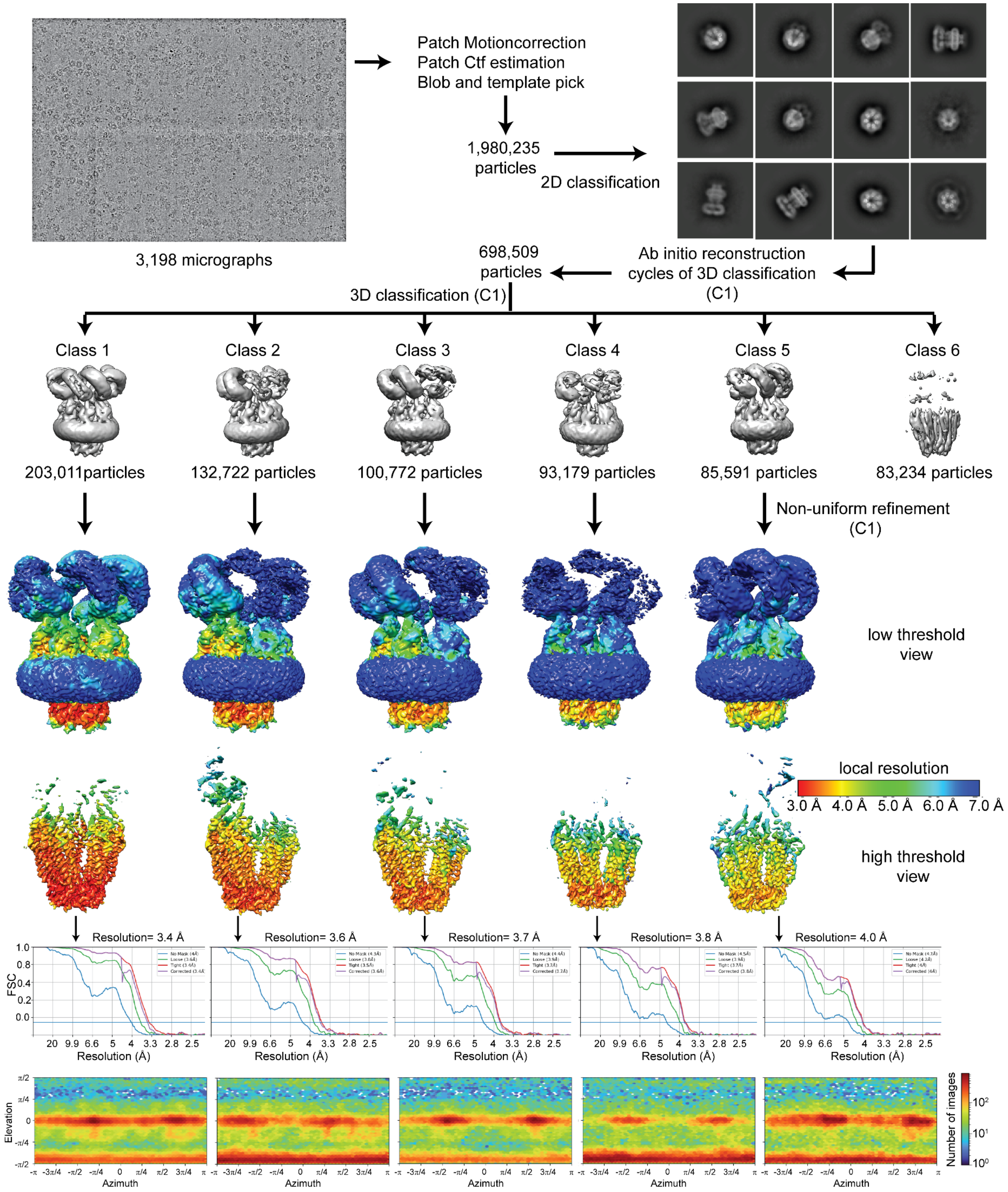


**Supplementary Fig. 2: Cryo-EM analysis of 8C-8A(IL1$^{25}$).** Flowchart detailing the particle selection and refinement procedure to obtain the cryo-EM maps of 8C-8A(IL1$^{25}$). Local resolution maps are shown on maps viewed at two different thresholds in Chimera. FSC curves and angular distributions are shown for the refined classes.

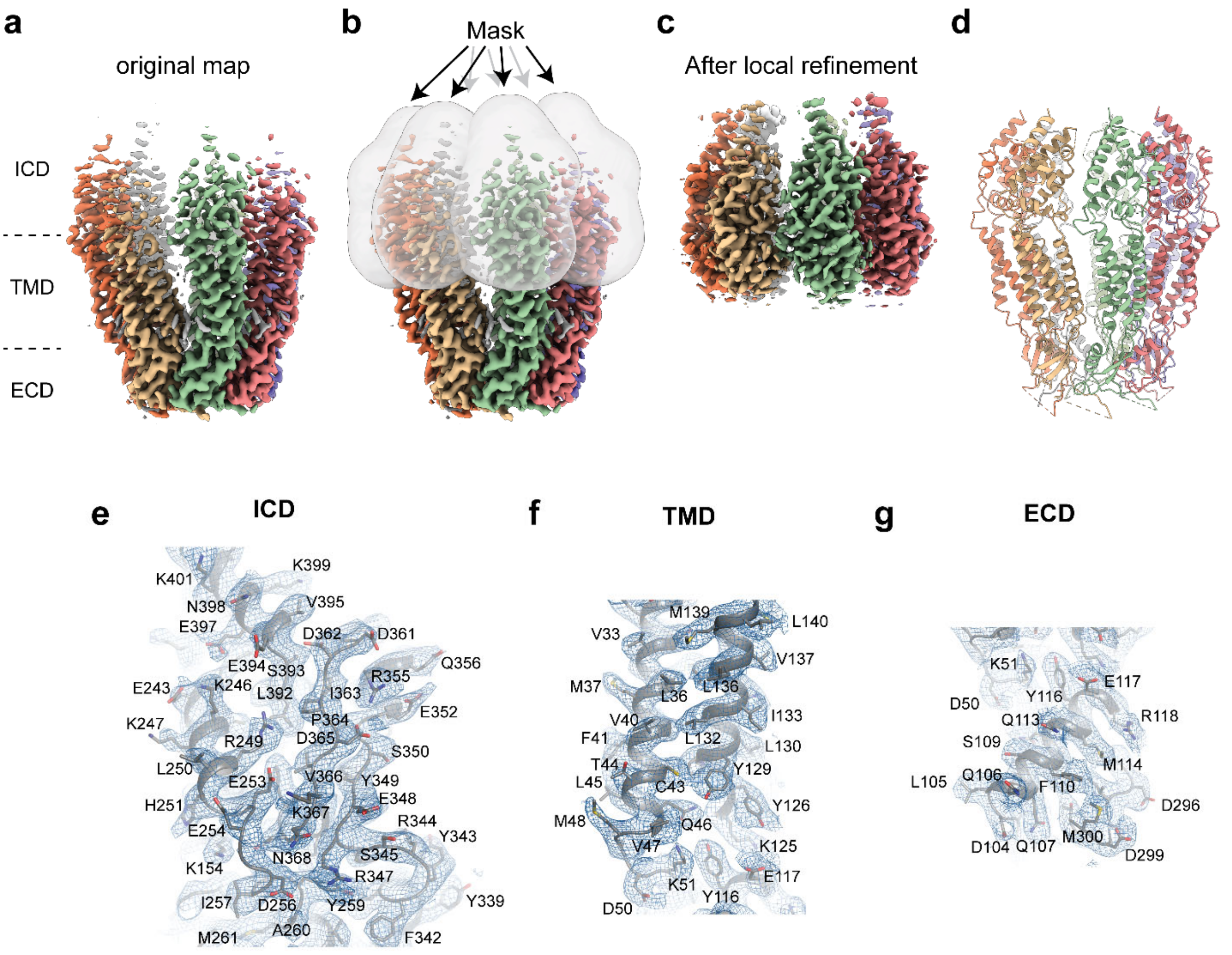


**Supplementary Fig. 3: Cryo-EM analysis of 8C-8A(IL1$^{25}$).** **a**) Cryo-EM map of 8C-8A(IL1$^{25}$) colored by individual protomers. **b**) Masks used for local refinement of individual protomers are shown as transparent grey surfaces. Each mask was used separately for local refinement. **c**) The cryo-EM maps after local refinement for each protomer are aligned onto the original map. **d**) Ribbon representation of the 8C-8A(IL1$^{25}$) structure. **e-f**) Representative images of the cryo-EM density and the modeled structure covering parts of ICD (**e**), TMD (**f**), and ECD (**g**). Map in panel e is obtained after local refinement, and maps in panels f and g are the original maps.

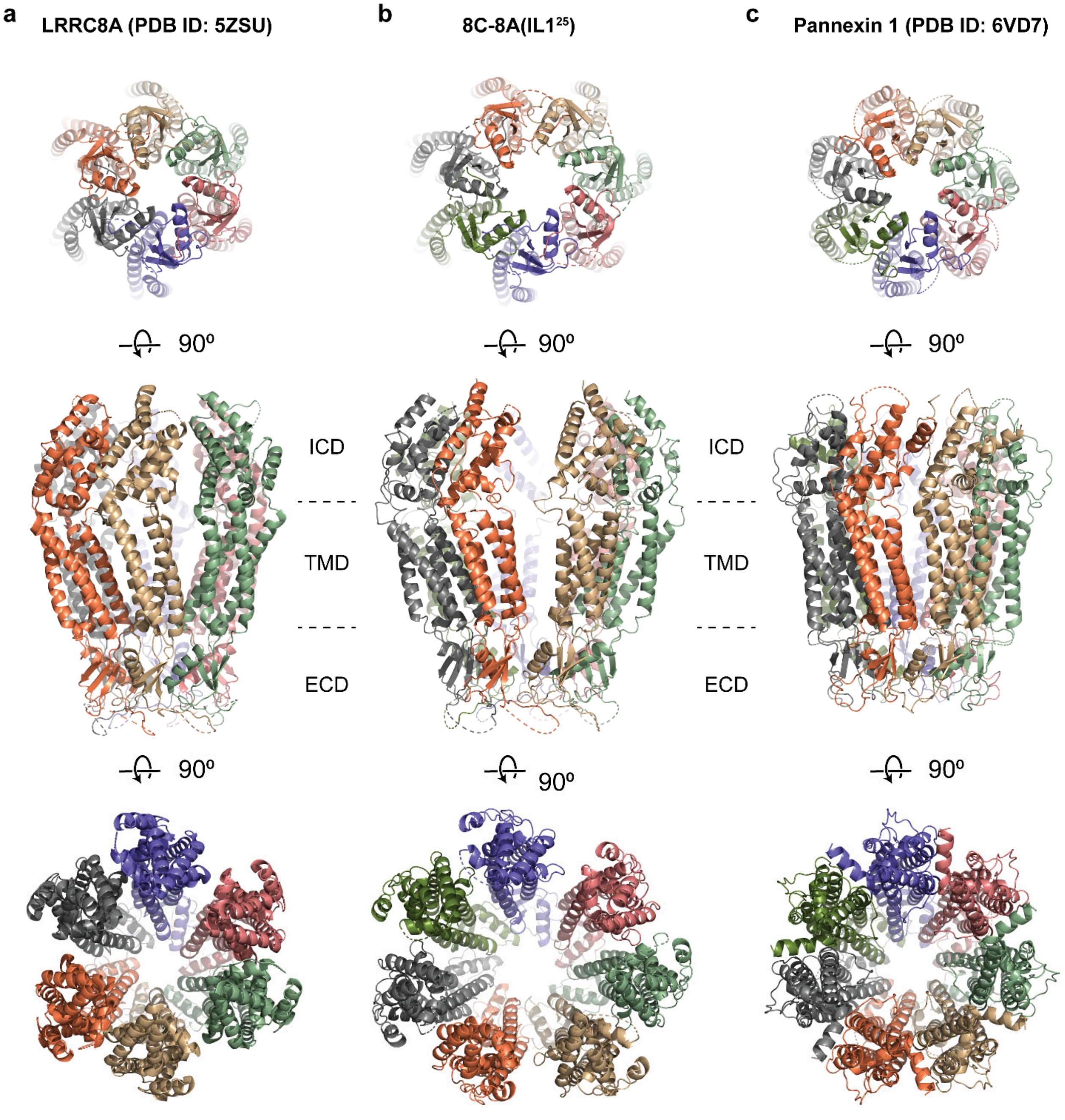


**Supplementary Fig. 4: Structural comparison of 8C-8A(IL1$^{25}$) to LRRC8A and pannexins. a-c)** Ribbon representations of LRRC8A (PBD ID: 5ZSU) (**a**), 8C-8A(IL1$^{25}$) (**b**), and Pannexin 1 (PDB ID: 6VD7) (**c**) viewed from the extracellular space (top), through the membrane plane (middle) and from the cytoplasm (bottom). LRRDs for 8C-8A(IL1$^{25}$) and LRRC8A were not shown. Each subunit is colored differently.

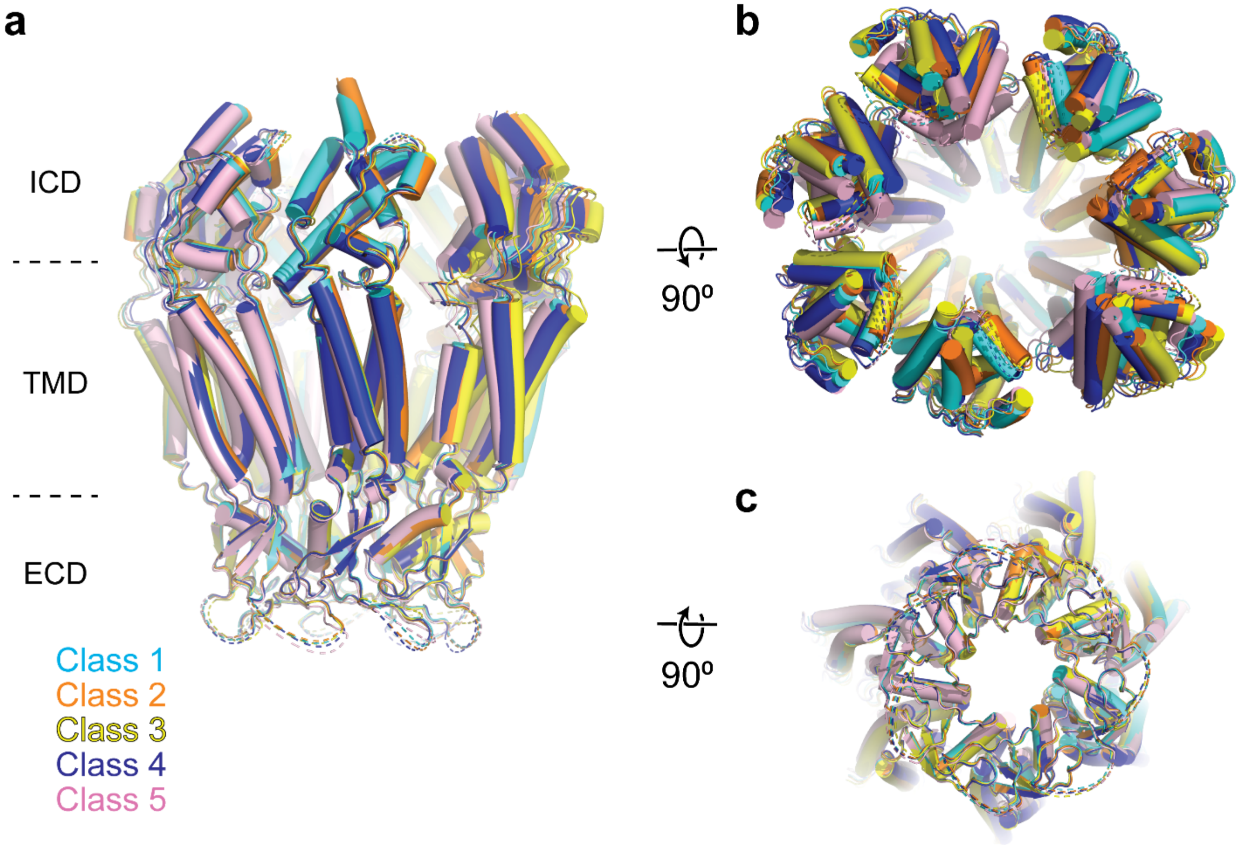


**Supplementary Fig. 5: Comparison of 8C-8A(IL1$^{25}$) structures. a-c)** Overlay of 8C-8A(IL1$^{25}$) structures in different classes viewed through the membrane plane (**a**), from the cytoplasm (**b**) and from the extracellular space (**c**). Structures are aligned on their ECDs and colored as indicated in the figure.

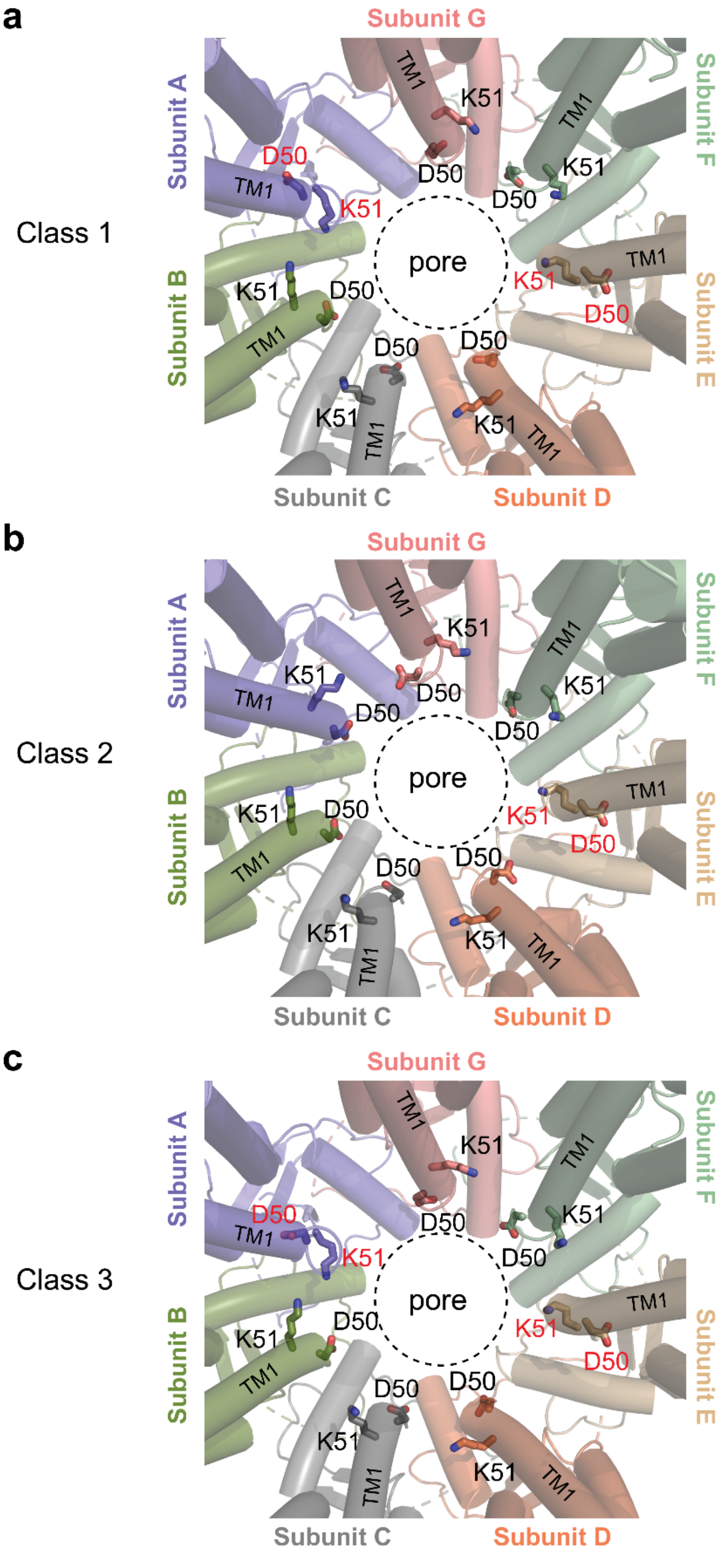


**Supplementary Fig. 6. Comparison of the D40-K51 loop among different classes.** Close-up view of the pore around the residues D50 and K51, which are shown as sticks. The residues that adopt different conformation compared to others are labeled in red. The dashed circle indicates the pore-lining surface.

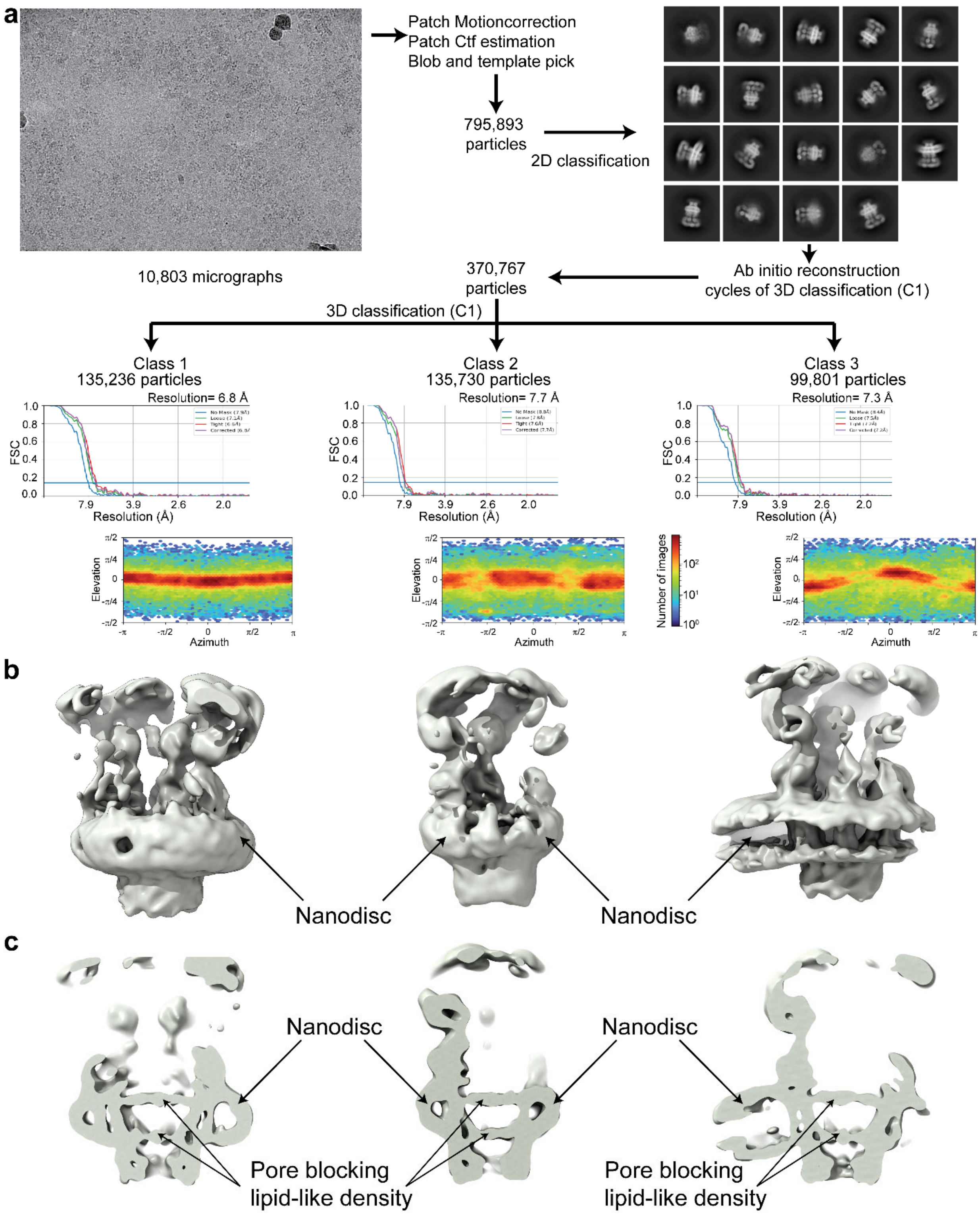


**Supplementary Fig. 7. Cryo-EM analysis of 8C-8A(IL1$^{25}$) reconstituted in nanodiscs. a)** Flowchart detailing the particle selection and refinement procedure to obtain the cryo-EM maps of the 8C-8A(IL1$^{25}$) nanodiscs. FSC curves and angular distributions are shown. **b-c**) Full (**b**) and sliced (**c**) views of the cryo-EM maps of the 8C-8A(IL1$^{25}$) nanodiscs.

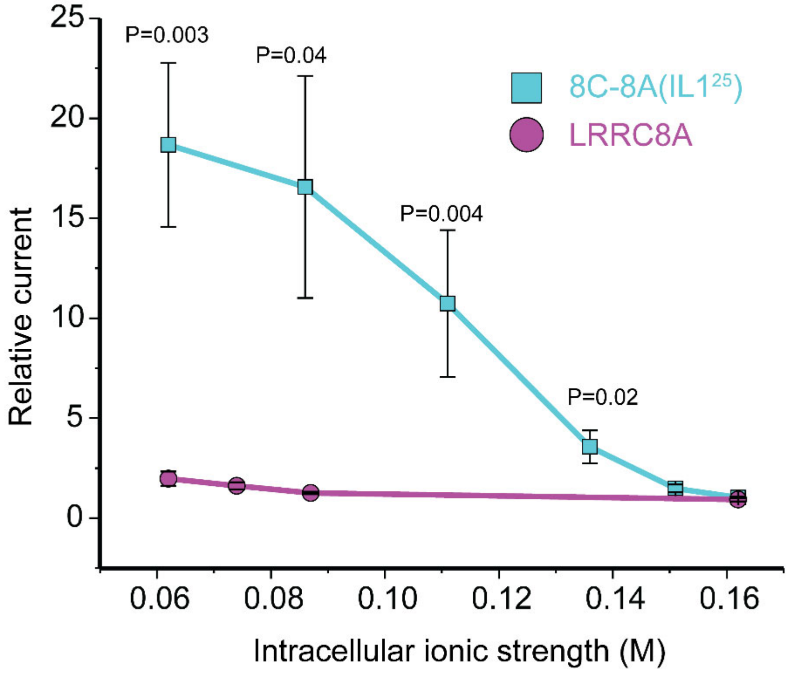


**Supplementary Fig. 8: Effect of intracellular ionic strength on activation of LRRC8A and the 8C-8A($IL1^{25}$) chimeric channel.** Relative current was quantified as the fold change in current measured immediately after whole cell access was achieved and currents measured 120 sec later. As we have shown previously[1], LRRC8A currents activate only very slowly when intracellular ionic strength is reduced from a normal level of 0.162 M to 0.062 M. However, no statistically significant (P>0.9) current activation is detected 120 sec after whole cell access even at the lowest intracellular ionic strength of 0.062 M. In striking contrast, significant (P=0.02) 8C-8A($IL1^{25}$) current is detected at an intracellular ionic strength of 0.136 M. Reducing ionic strength further leads to more rapid current activation and larger current amplitudes measured at 120 sec. The effect of ionic strength on the rate of (8C-8A($IL1^{25}$) current activation is similar to what we have reported previously for native VRAC/LRRC8 channels[2]. Current amplitude was quantified at +60 mV above the measured reversal potential. Values are means ± SEM (N=5-8).

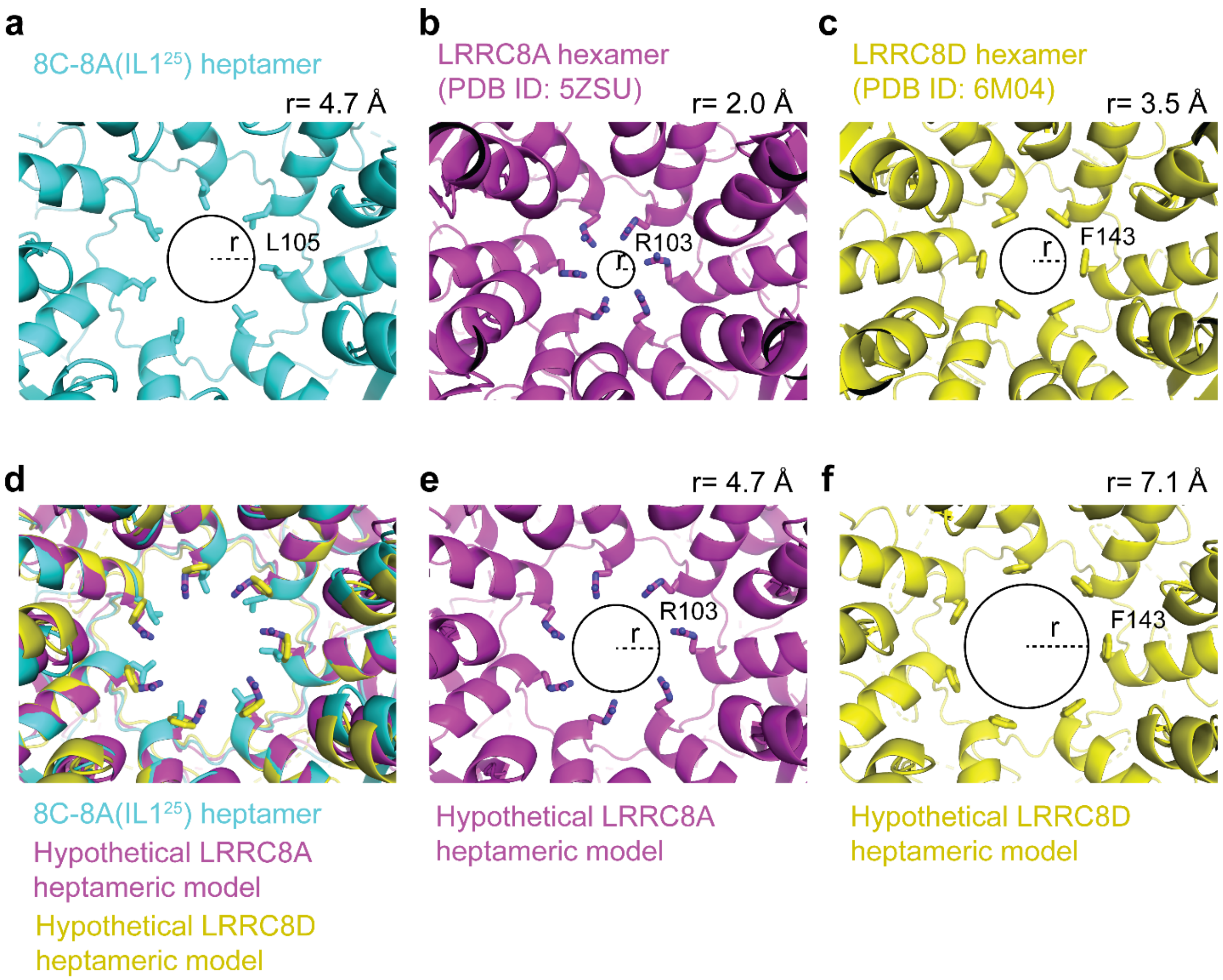


**Supplementary Fig. 9: Heptameric oligomerization leads to larger pore sizes. a-c**) Close-up cytoplasmic view of the narrowest constriction sites of 8C-8A(IL1[25]) (**a**), LRRC8A (PDB ID: 5ZSU) (**b**), and LRRC8D (PDB ID: 6M04) (**c**). **d**) The heptameric LRRC8A and LRRC8D models were obtained by overlaying one of the LRRC8A or LRRC8D protomers on the seven protomers of 8C-8A(IL1[25]) heptamer by aligning the ECDs. e-f) Pores of hypothetical LRRC8A (e) and LRRC8D (f) heptamers. Residues forming the constriction are shown as sticks. The circles and the dashed lines represent the solvent accessible pore size and radius, respectively. The radii of the pores were calculated using the software HOLE[3] and shown on top of each panel.

### Supplementary Table 1: Cryo-EM data collection, refinement, and validation statistics

| **Data collection and processing** | | | | | |
|---|---|---|---|---|---|
| Microscope | FEI Krios G3i microscope | | | | |
| Detector | Gatan K3 direct electron camera | | | | |
| Nominal magnification | 81,000 x | | | | |
| Voltage (kV) | 300 | | | | |
| Electron exposure (e/Å$^2$) | 54 | | | | |
| Defocus range (µm) | -0.8 to -1.5 | | | | |
| Pixel size (Å) | 1.1 | | | | |
| Number of Micrographs | 3,198 | | | | |
| Particles images (no.) | 846,122 | | | | |
| Conformational state | **Class 1** | **Class 2** | **Class 3** | **Class 4** | **Class 5** |
| Symmetry imposed | C1 | C1 | C1 | C1 | C1 |
| Final particles images (no.) | 203,011 | 132,722 | 100,772 | 93,179 | 85,591 |
| Map resolution (Å) (FSC threshold=0.143) | 3.4 | 3.6 | 3.7 | 3.8 | 4.0 |
| **Refinement** | | | | | |
| Model resolution (Å) (original map, FSC threshold=0.5) | 3.6 | 3.9 | 3.9 | 4.2 | 4.4 |
| B-factor used for map sharpening (Å$^2$) | -102.0 | -88.2 | -84.7 | -70.5 | -83 |
| ***Model composition*** | | | | | |
| Non-hydrogen atoms | 17,499 | 17,499 | 17,465 | 10,435 | 10,435 |
| Protein residues | 2,101 | 2,101 | 2,097 | 2,101 | 2,101 |
| ***Mean B factors (Å$^2$)*** | | | | | |
| Protein | 123.52 | 89.5 | 110.6 | 248.8 | 247.6 |
| ***R.m.s. deviations*** | | | | | |
| Bond lengths (Å) | 0.003 | 0.002 | 0.003 | 0.002 | 0.004 |
| Bond angles (°) | 0.563 | 0.525 | 0.553 | 0.516 | 0.783 |
| ***Molprobity score*** | 1.72 | 1.69 | 1.75 | 1.37 | 1.49 |
| ***Clash score*** | 5.09 | 4.95 | 5.94 | 1.14 | 1.54 |
| ***Poor rotamers (%)*** | 0.0 | 0.0 | 0.0 | 0.0 | 0.0 |
| Favored (%) | 92.9 | 93.5 | 93.6 | 89.95 | 88.3 |
| Allowed (%) | 7.1 | 6.5 | 6.4 | 10.0 | 11.5 |
| Disallowed (%) | 0 | 0 | 0 | 0.05 | 0.15 |
